# Emergence of a Novel Bacteriophage of Toxigenic *Vibrio cholerae* in Haiti

**DOI:** 10.64898/2026.11.15.766889

**Authors:** Meer Taifur Alam, Andrea Rebolledo-Viveros, S. Nazmus Sakib, Francklin Brice Soung Iballa, J. Glenn Morris, Andrew Camilli, Afsar Ali

**Affiliations:** Emerging Pathogens Institute, University of Florida, Gainesville, FL, USA; Department of Environmental and Global Health, College of Public Health and Health Professions, University of Florida, Gainesville, FL, USA; Department of Molecular Biology and Microbiology, Graduate School of Biomedical Sciences; School of Medicine, Tufts University, Boston, MA, USA

## Abstract

Toxigenic strains of *Vibrio cholerae* O1 are facultative pathogens that persist in estuaries and cause the secretory diarrheal disease cholera. Virulent bacteriophages (phages) can modulate pathogen density in both humans and aquatic reservoirs, thereby reducing cholera transmission. We report a novel phage, MAA01, isolated from cholera stool and environmental water samples collected in Haiti. The MAA01 genome is double-stranded DNA (31,507 base pairs) and contains 47 genes. Based on transmission electron microscopy morphology and phylogenetic analysis, we classify the novel phage as a member of the Caudoviricetes class. Its genome content is consistent with a temperate lifestyle, yet we observed a virulent phenotype in all strains tested, including toxigenic and non-toxigenic environmental O1 serogroup strains. Sequence analysis of spontaneous escape mutants of *V. cholerae* identified the O1 lipopolysaccharide antigen as the phage receptor. Using an infant mouse model of intestinal colonization, we showed that MAA01 can reduce *V. cholerae* load by 4-fold, indicating it can prey on its host in the small intestine.

**IMPORTANCE:** Cholera remains a major cause of diarrheal disease, and virulent phages that prey on *V. cholerae* can shape its abundance in patients and in aquatic reservoirs, thereby influencing transmission. Until now, ICP2 was the only virulent phage reported in Haiti, where a cholera epidemic began in 2010. Here we describe MAA01, a novel phage recovered from both cholera patient stools and surface waters in Haiti. MAA01 uses the O1 lipopolysaccharide antigen as its receptor, restricting its host range to serogroup O1, and it reduces *V. cholerae* burden in the infant mouse small intestine. These findings expand the known diversity of *V. cholerae* phages, identify a candidate for phage-based prophylaxis or therapy, and argue for broader phage surveillance in cholera-endemic settings.

## INTRODUCTION

The nation of Haiti was cholera-free for over a century until October 2010, when peacekeeping troops, providing humanitarian support to Haitians devastated by a massive Earthquake in January 2010, unknowingly brought in cholera (1–3). Just under 820,000 cholera cases, with nearly 10,000 deaths, were recorded in Haiti between 2010 and 2018 (4). Although the epidemic was thought to have subsided in 2018, cholera cases reappeared in 2022, with evidence of ongoing endemic disease continuing through 2026 (5–7). Cholera is caused by toxigenic strains of *V. cholerae* serogroup O1, whose genome harbors the filamentous temperate phage CTXФ, which encodes cholera toxin and promotes profuse secretory diarrhea (8). We and others previously demonstrated that a toxigenic *V. cholerae* O1 strain with a high degree of genomic similarity to Asian cholera strains was introduced into Haiti in 2010 and was responsible for the Haitian cholera epidemic (1, 3).

*V. cholerae* is a natural resident of aquatic reservoirs, and humans acquire the disease by consuming contaminated food or water (9–13). *V. cholerae* has two lifestyles: a transient one in a human host and a permanent one in aquatic reservoirs (11, 13). To establish cholera endemicity in a new region, as in Haiti, *V. cholerae* must adapt to and evolve in the face of stressful conditions in both human hosts and aquatic reservoirs, including host immunity, nutrient limitation in aquatic reservoirs, variables such as temperature and salinity, and predation by protists and phages (14–19). *V. cholerae* phages have been shown to influence cholera infection dynamics and persistence in aquatic reservoirs by limiting and/or altering transmission dynamics (20–22).

Several cholera-endemic countries have reported virulent *V. cholerae* phages. They are generally isolated from cholera stool samples but are sometimes also isolated from sewage and bodies of water (21–25). Several recent studies demonstrated that the prevalence and occurrence of *V. cholerae* phages vary geographically. For example, a robust analysis of cholera stool samples collected over a ten-year period in Bangladesh showed the coexistence of three virulent Caudoviricetes class phages: Mohonavirus-genus phage ICP1, Icepovirus-genus phage ICP2, and Chatterjeevirus-genus phage ICP3 (21). In contrast, we recently reported ICP1, but not ICP2 or ICP3, in the Democratic Republic of Congo (5). In another example, we reported ICP2, but not ICP1 or ICP3, after examining cholera stool samples in Haiti (24). The Haiti ICP2 isolate is genetically highly related to contemporary Asian ICP2 isolates (24) and was therefore likely introduced into Haiti in 2010 along with toxigenic *V. cholerae*. *V. cholerae* phages have also been isolated and characterized from other parts of the world (25, 26). ICP1, ICP2, and ICP3 prey on *V. cholerae* in the human small intestine, thereby imposing selective pressure (21, 22, 24, 27, 28) and reducing *V. cholerae* load in animal models of cholera (29).

Because of the unique history of cholera in Haiti, namely its reintroduction in 2010 after more than a century of being cholera-free, plus its geographical isolation from cholera-endemic countries, we were interested in knowing whether other *V. cholerae* phages besides ICP2 were present. Therefore, we conducted an extensive search for additional phages. Here we report the isolation and characterization of a novel phage from both cholera stool samples and aquatic reservoirs in Haiti, which we have designated phage MAA01 (<u>M</u>eer-<u>A</u>fsar-<u>A</u>li). MAA01 is highly specific to *V. cholerae* O1 hosts because it uses the lipopolysaccharide O1-antigen as its receptor. We report MAA01 predation parameters *in vitro* and show that it can prey on *V. cholerae* in the small intestine and reduce bacterial load in a murine model of intestinal colonization.

## RESULTS AND DISCUSSION

### Prevalence of MAA01 phage in clinical and environmental samples isolated in Haiti

We initially screened 75 stool samples from confirmed cholera patients and 30 freshwater, estuarine, and sewage surface water samples collected during 2014-2017 for phages that plaque *V. cholerae* O1. We collected clinical samples at cholera treatment centers (CTCs) and clinics across Haiti and environmental samples from nearby sources (**Fig. 1**). We detected *V. cholerae* O1 in stool samples by standard enrichment followed by streaking on an agar medium selective for *V. cholerae*. Phages in environmental samples were enriched by allowing multiplication on two *V. cholerae* O1 strains before assaying for phage plaques on soft agar overlays. PCR was done on individual plaques to detect ICP1, ICP2, or ICP3. We detected ICP2 but not ICP1 or ICP3, consistent with a previous report (24). We subjected single, non-ICP2 plaques isolated from five clinical and three environmental samples to whole-genome sequencing. All eight sequences were identical, revealing a novel phage, designated MAA01.

**Figure 1.**
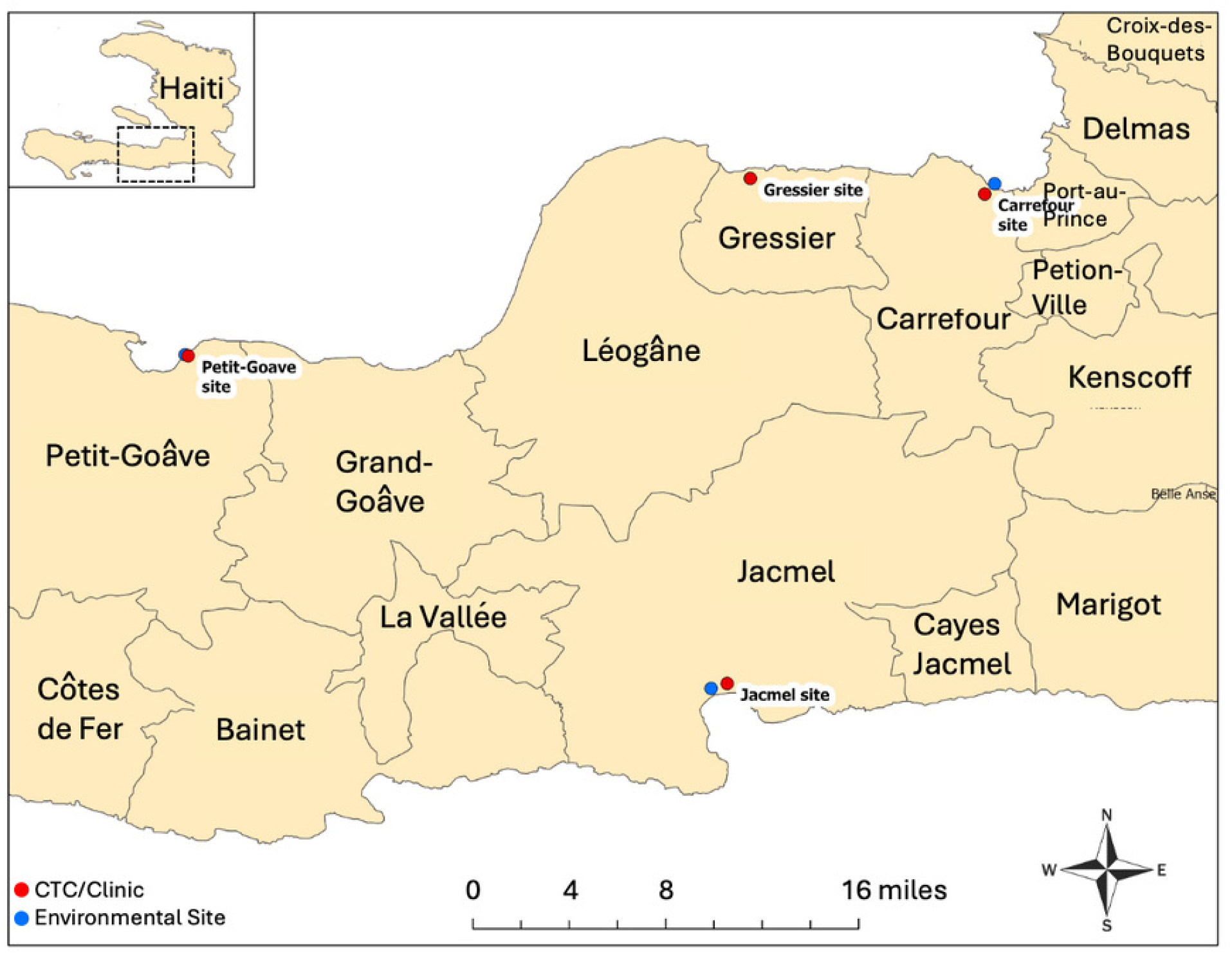
Map showing sites of sample collection. Locations at which we collected suspected cholera stool samples and environmental water samples in Haiti for isolation of both *V. cholerae* O1 strains and phage are shown. Solid red and blue filled circles indicate cholera treatment centers (CTC)/clinics and environmental sites, respectively.

We used PCR with MAA01-specific primers to screen phages isolated from plaques from all stool and environmental samples in our 2014-2017 collection. In subsequent studies, we also screened plaque isolates from 19 stool samples from cholera patients collected between 2017 and 2020, 50 stool samples collected during that same period from persons culture-negative for *V. cholerae*, and 47 stool samples from cholera patients from 2022, when cholera recurred in Haiti. Results are shown in Table 1. In the initial 2014-2017 group of clinical and environmental samples, 25 (33.3%) of 75 clinical samples were positive for MAA01 (**Tables 1, S1**), as were 7 of the 30 environmental samples screened (**Tables 1, S2**). None of the environmental samples tested were positive for toxigenic *V. cholerae* O1; the absence of *V. cholerae* in the environmental samples is not unusual, as only a few percent of environmental samples in Haiti yield *V. cholerae* (30, 31). The frequency of MAA01 identification in clinical isolates was lower in subsequent years, with plaques from 1 of 19 clinical samples from 2017-2020 and 2 of 47 clinical samples from 2022 positive for MAA01. Taken together, our data indicate that the MAA01 phage is present both in cholera patient stool samples and environmental water samples in Haiti. Notably, MAA01 was not detected in any stool or environmental samples collected before March 11, 2015, but then appeared in stool samples collected in the coastal city of Jacmel (**Table S1**), with subsequent spread to other sites in Haiti. This pattern is consistent with the hypothesis that MAA01 jumped from the coastal aquatic environment in the Jacmel region into humans, followed by spread together with stool-shed *V. cholerae*. Phylogenetic studies have implicated environmental *V. cholerae* O1 Ogawa strains from Jacmel as the source for the recurrent cholera epidemic which began in 2022 (5); it is plausible that MAA01 phage were “fellow travelers” in this movement.

**Table 1.**
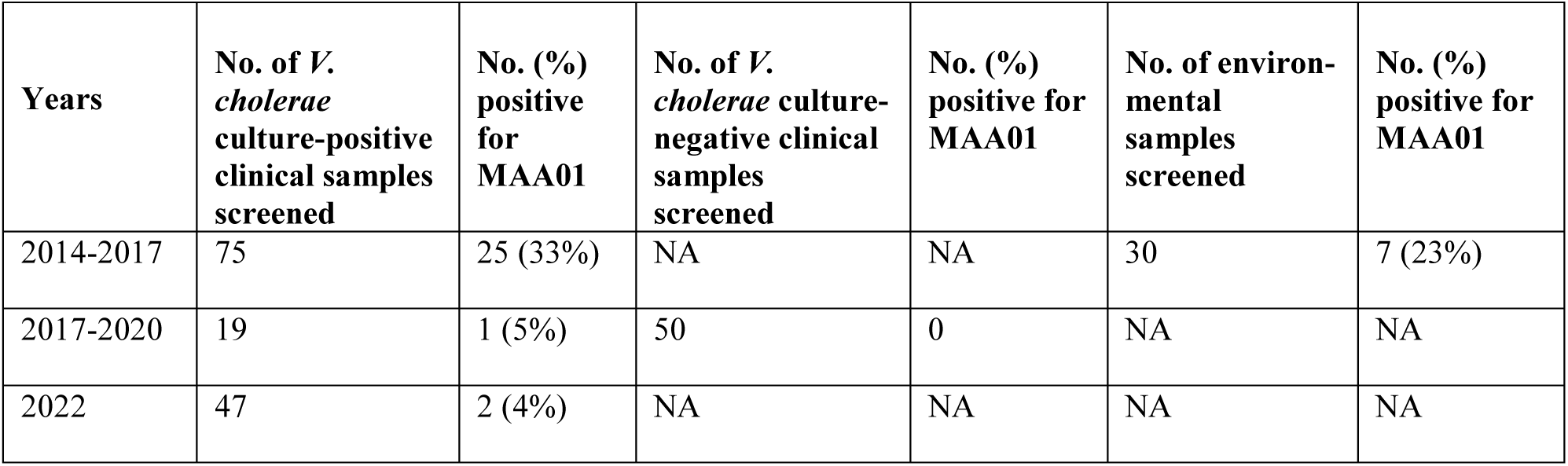
Prevalence of MAA01 phage in cholera stool and environmental samples collected.

| **Years** | **No. of *V. cholerae* culture-positive clinical samples screened** | **No. (%) positive for MAA01** | **No. of *V. cholerae* culture-negative clinical samples screened** | **No. (%) positive for MAA01** | **No. of environ-mental samples screened** | **No. (%) positive for MAA01** |
|---|---|---|---|---|---|---|
| 2014-2017 | 75 | 25 (33%) | NA | NA | 30 | 7 (23%) |
| 2017-2020 | 19 | 1 (5%) | 50 | 0 | NA | NA |
| 2022 | 47 | 2 (4%) | NA | NA | NA | NA |

### Morphology of MAA01 phage

To determine MAA01 morphology, we examined a concentrated, pure phage sample by transmission electron microscopy. The phage had a hexagonal head (∼58 nm in diameter) and a short tail (∼16 nm long) (**Fig. 2**). Based on morphology, MAA01 belongs to the Caudoviricetes class of phages.

**Figure 2.**
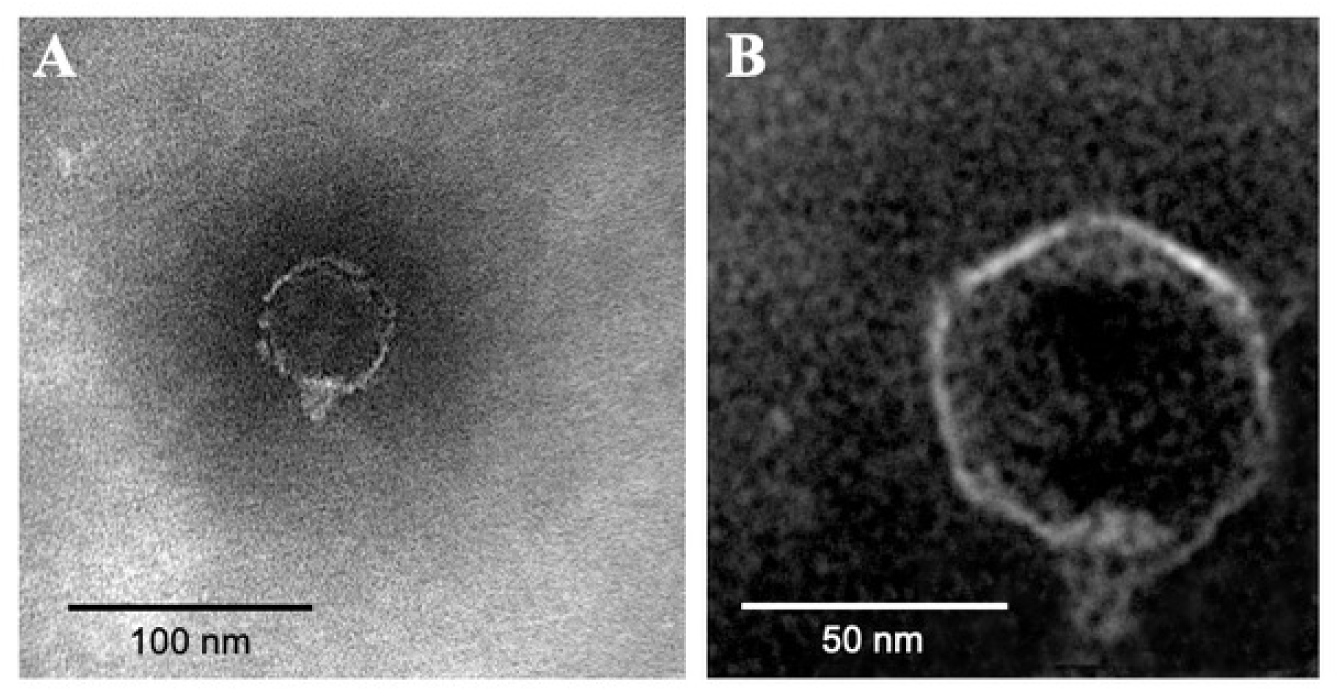
Micrograph of MAA01 phage detected by transmission electron microscopy. The phage was negatively stained with 1% uranyl acetate and viewed by transmission electron microscopy at 50,000х magnification. The micrographs (**A, B**) collectively show an icosahedral head (∼58 nm in diameter) and a short (∼16 nm) tail.

### MAA01 phage genome

The MAA01 phage genome is a double-stranded DNA molecule totaling 31,507 nucleotides, with a 47% G/C content. The complete MAA01 genome sequence was deposited in GenBank (Accession No. QB327871). The genome contains 47 protein-encoding genes as determined by PROKKA and ORF Finder software (32) (**Fig. 3**, **Table S3**). Of the 47 genes, 27 encode hypothetical proteins, as none show similarity to any known protein in the NCBI protein database. The remaining 20 gene products fall into three major phage functional categories: head/tail-related proteins (n=4), capsid proteins (n=2), and DNA replication/metabolism proteins (n=14). MAA01 contains two gene products found in temperate phages, an integrase and a transcriptional repressor. Accordingly, the phage analysis program PhageScope (33) classified MAA01 as a temperate phage. However, when we tested a variety of environmental and clinical O1 serogroup strains isolated from different countries over 50 years, we observed only clear plaques rather than cloudy plaques, suggesting that MAA01 could not lysogenize (**Table 2**, **Table S4)**. Furthermore, after passage (predation) of MAA01 on two of these strains, Haiti strain HC1037 and Bahrain strain E7946, in broth culture, we failed to obtain colonies resistant to superinfection (i.e., lysogens). This could be because the *V. cholerae* strains tested lack the MAA01 attachment site (attB), or because the putative integrase is not functional, although other reasons may apply. It is unlikely that the integrase is nonfunctional, given that the protein is full length compared to related phage integrases and contains all highly conserved active-site residues of tyrosine-type recombinase/integrase enzymes.

**Figure 3.**
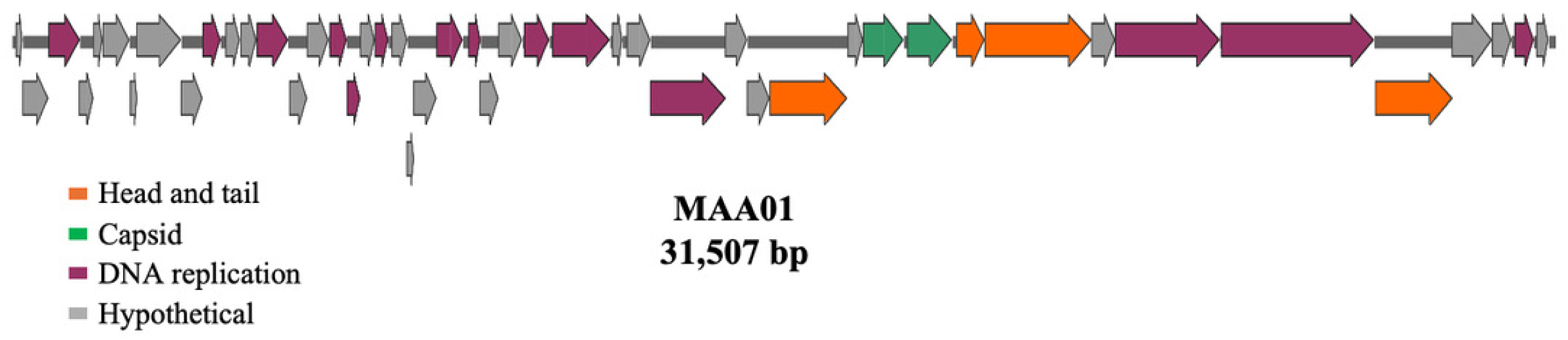
Genetic map of MAA01 phage. The MAA01 phage genome encodes 31,507 base pairs and 47 predicted protein-coding genes. The transcription direction of each gene is shown by arrows (both in upper and lower rows), and each protein product’s potential function is indicated in a color-coded format: head- and tail-related proteins (orange), capsid protein (green), DNA replication and related proteins (magenta), and hypothetical proteins (gray). The potential gene function was determined by comparing each gene to that of existing nucleotide and/or protein sequences available in the GenBank database (NCBI).

**Table 2.** Host range for MAA01 phage.

| **Strains with description** | **Source/year of isolation** | **Susceptibility to MAA01 phage** |
|---|---|---|
| *V*. *cholerae* O1 N16961-Inaba (El Tor) | Bangladesh, 1975 | S |
| *V*. *cholerae* O1 O395-Ogawa (Classical) | India, 1965 | S |
| *V*. *cholerae* NRT 36 | Japan, 1984 | R |
| *V*. *cholerae* O139 (1837) | Peru, 1994 | R |
| *E*. *coli* DH5alpha | Lab strain, USA | R |
| *V*. *cholerae* O139, MO10 | India, 1992 | R |
| *V*. *cholerae* non-O1, non-O139 | Chilli | R |
| *V*. *cholerae* non-O1, non-O139 CR-03424 | USA, 2003 | R |
| *V*. *cholerae* non-O1 1587 | Peru, 1994 | R |
| *V*. *cholerae* non O1(Polz) | USA, 2006 | R |
| *V*. *cholerae* non O1 strain 062478 (Polz) | USA, 2006 | R |
| *V*. *parahaemolyticus* CAB2 tdh-, trh-, exsA- | Laboratory collection, USA | R |
| *V*. *parahaemolyticus* 06071 TB3 | Laboratory collection, USA | R |
| *V*. *harveyi* BB120 | Laboratory collection, USA | R |
| *V*. *splendidus* ZE90 (Polz) | Laboratory collection, USA | R |
| *V*. *splendidus* 12-B01(Polz) | Laboratory collection, USA | R |

### Identifying the receptor for MAA01 phage

To determine the host receptor utilized by MAA01, we isolated multiple independent spontaneous MAA01-resistant escape mutants of *V. cholerae* O1 strains E7946 and HC1037. E7946 is a clinical strain from Bahrain isolated in 1978 and represents an early (wave 1) 7^th^ pandemic strain, whereas HC1037 is a clinical strain from Haiti isolated in 2014 and represents a recent (wave 3) 7^th^ pandemic strain. We compared whole-genome sequences of escape mutants to those of the wild-type parent strains. All mutations mapped to lipopolysaccharide O1-antigen biosynthetic genes *gmD*, *wbeG*, *wbeL*, or *wbeN* (**Fig. 4**, **Tables S5, S6**). These data indicate that the *V. cholerae* O1 antigen is the receptor for the MAA01 phage. This is the same receptor used by phages ICP1 and ICP3 (21).

**Figure 4.**
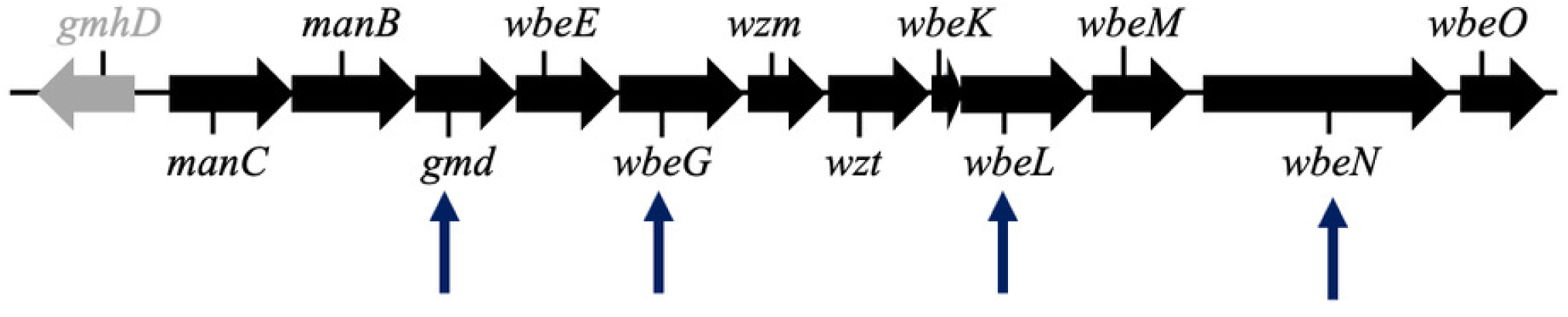
Genetic map of the O1 antigen biosynthetic gene cluster showing mutated genes. The annotated genes and their orientation are indicated by block arrows (drawn to scale). Genes in black are O1 antigen biosynthetic genes, while *gmhD* (gray) is a flanking gene. Spontaneous MAA01-resistant escape mutations mapped to the genes indicated by the vertical arrows.

### Phylogenetic analysis

To determine whether MAA01 is a novel phage, we compared its genome with complete phage genomes, including known *V. cholerae* phages deposited in NCBI’s GenBank. To determine the phylogeny of MAA01, we used Virfam software to run a Hidden Markov model search (34) with five conserved proteins: the major capsid, portal, terminase large subunit, adaptor, and head-closure proteins, with those of phages deposited in the Aclame database (1,071 phage genomes). The resulting confidence probabilities classified MAA01 as a tailed dsDNA phage within the Caudoviricetes class. Despite the high HMM search confidence probabilities (100%) for all five proteins, the sequence identity of the closest protein in database matches did not exceed 28%, indicating that MAA01 was not closely related to any (**Table S7**). Additionally, we performed a phylogenetic analysis of MAA01 and 82 phages using a conserved gene product, the terminase large subunit. MAA01 was distantly related to all but two marine phages of *Vibrio splendidus* (**Fig. 5**). The two *V. splendidus* phages are syntenic and share 93% nucleotide identity, but are presently unclassified at the family, genus, and species levels. MAA01 is not syntenic with either phage and only shares 69% nucleotide identity with each. Based on these data, we conclude that MAA01 is a novel *V. cholerae* O1-specific phage in the Caudoviricetes class, but it cannot be classified further.

**Figure 5.**
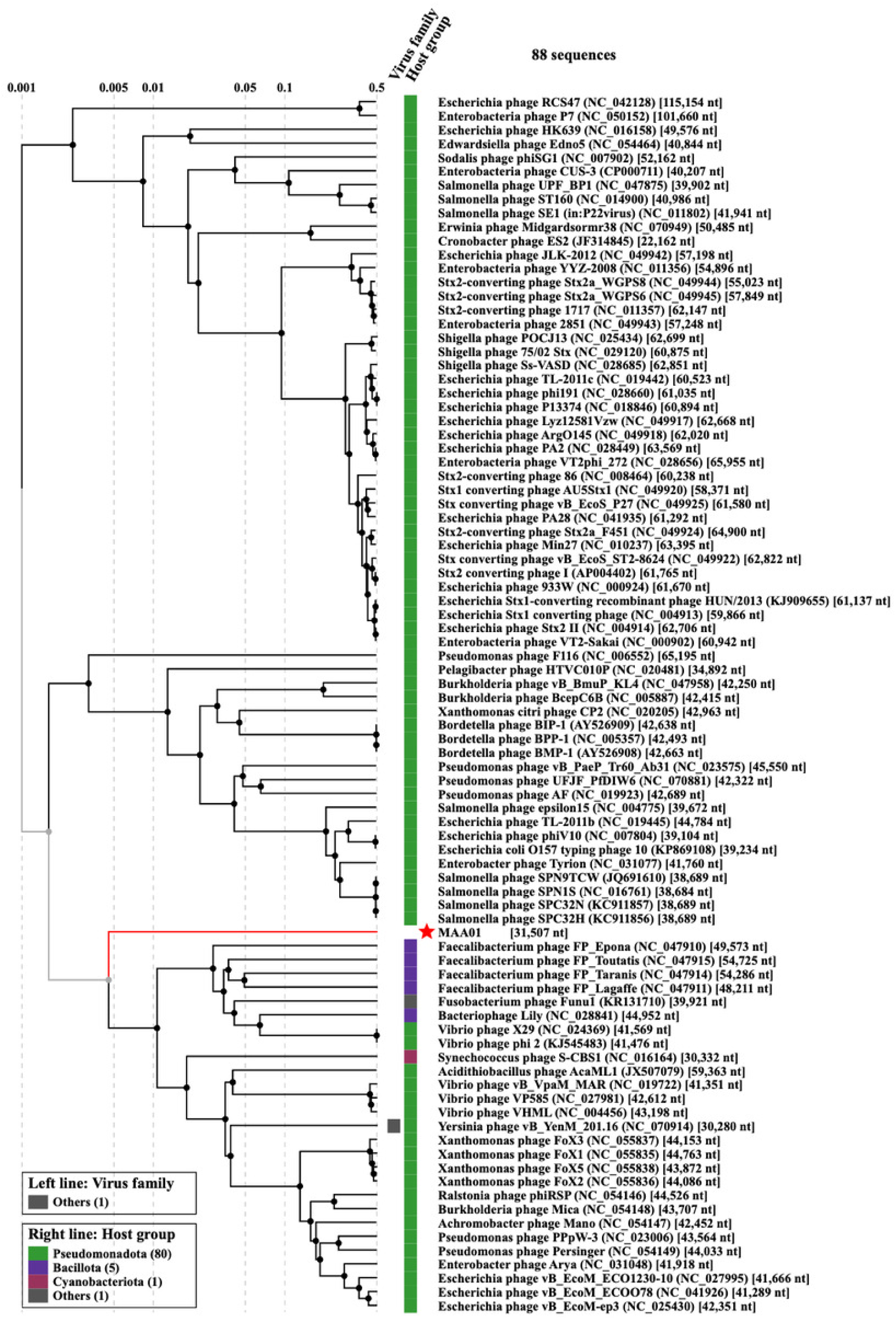
Phylogenetic tree for MAA01 phage. We constructed a phylogenetic tree of MAA01 using the neighbor-joining method and compared the amino acid sequence of the large terminase subunit of the MAA01 phage with that of 82 Caudoviricetes phages (obtained from GenBank). Different color groups indicate the DNA packaging strategy and phage type for each group. The black arrow indicates the novel MAA01 phage.

### Host range and multiplication parameters for MAA01 phage

To determine the host range of MAA01, we assayed plaque formation on a collection of O1, O139, and non-O1 *V. cholerae* as well as a variety of other *Vibrio* species (**Table 2**). MAA01 formed plaques only on *V. cholerae* O1 strains, consistent with the findings above that it requires the O1 antigen as its receptor. To determine phage adsorption time, eclipse time, burst time, and burst size, we used *V. cholerae* HC1037 as the host, as described in the Materials and Methods section. MAA01 was nearly 100% adsorbed onto its host by 9 min at 37 °C (**Fig. 6A**). The phage eclipse and burst times were both 50 min at 37 °C, and were 60 and 80 min at 25 °C, respectively (**Fig. 6B)**. The burst size was approximately 95 PFU at both temperatures (**Fig. 6C**).

**Figure 6.**
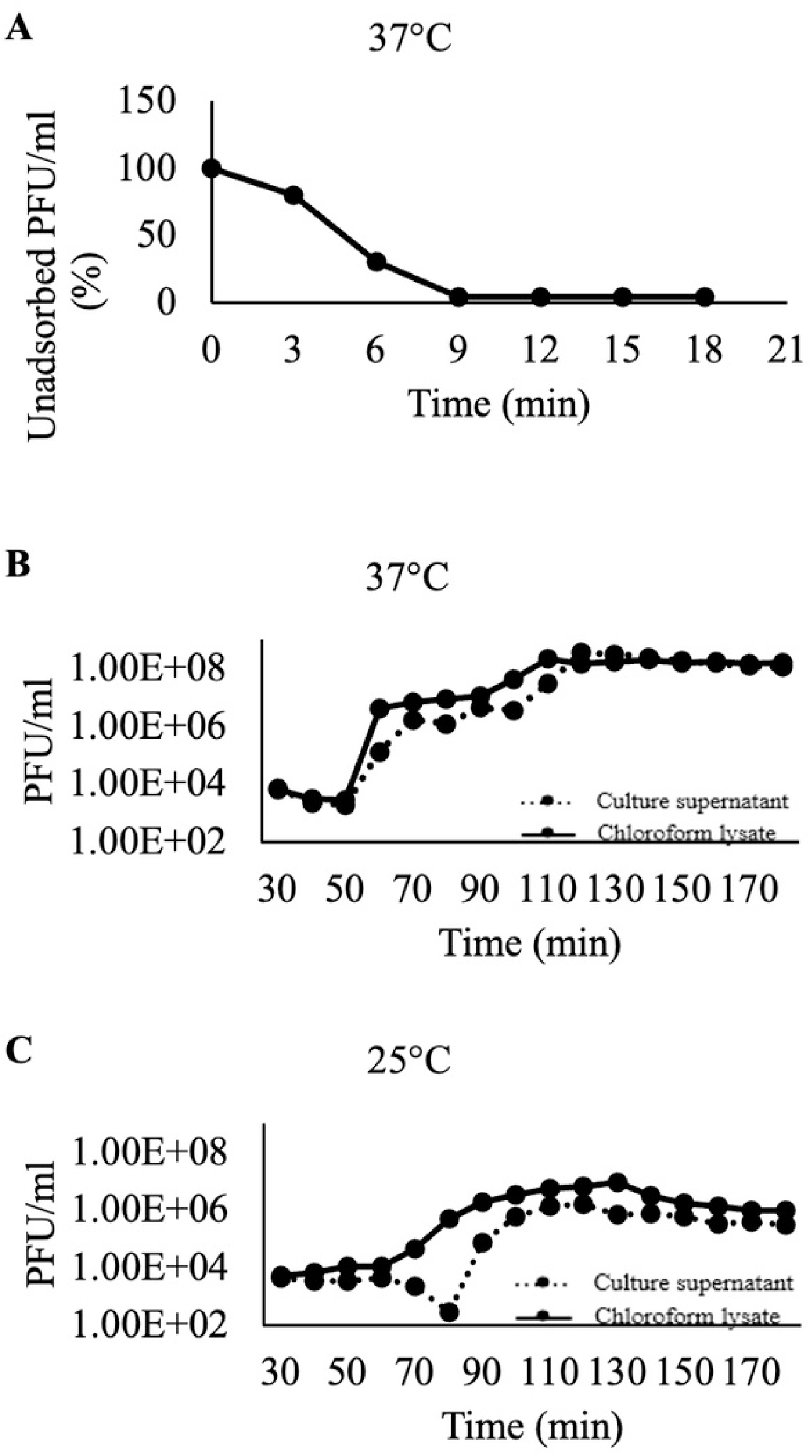
Measurement of MAA01 phage infection characteristics. The adsorption, eclipse time, burst time, and burst size upon exposure to *V. cholerae* strain HC1037 (a clinical toxigenic strain isolated from Haiti). We observed that MAA01 phage was nearly 100% adsorbed to its host by 9 min, with an eclipse time of ∼40 min (**A**). The burst time was 50 min (**B**) and 80 min (**C**) at 37 °C and 25 °C, respectively, and the burst size was ∼95 PFU at both temperatures.

### MAA01 phage is active in the murine small intestine

Our frequent isolation of MAA01 from stool samples of cholera patients suggested that it can prey on *V. cholerae* in the small intestine. To test this hypothesis, we assessed whether MAA01 could reduce *V. cholerae* burden in the small intestine of infant mice when administered prophylactically, as previously described for ICP1, ICP2, and ICP3 (29). Two groups of eight mice were administered MAA01 phage intragastrically and, 3.5 or 7 hours later, were challenged with *V. cholerae* O1 strain HC1037. A third group received no phage but was challenged with HC1037. After 21 hours of infection, we determined the *V. cholerae* load in the small intestine. MAA01 reduced the load of *V. cholerae* 4-fold in the 3.5 h prophylaxis group, but did not significantly reduce the load in the 7 h prophylaxis group (**Fig. 7**). These data show that MAA01 phage is active in predation within the small intestine, consistent with its frequent isolation from stool samples from cholera patients in Haiti. However, the observed 4-fold reduction in bacterial load in infant mice is far less than the 2- to 6-order-of-magnitude reduction reported for ICP1, ICP2, and ICP3 (27), suggesting MAA01 is less active in the small intestine.

**Figure 7.**
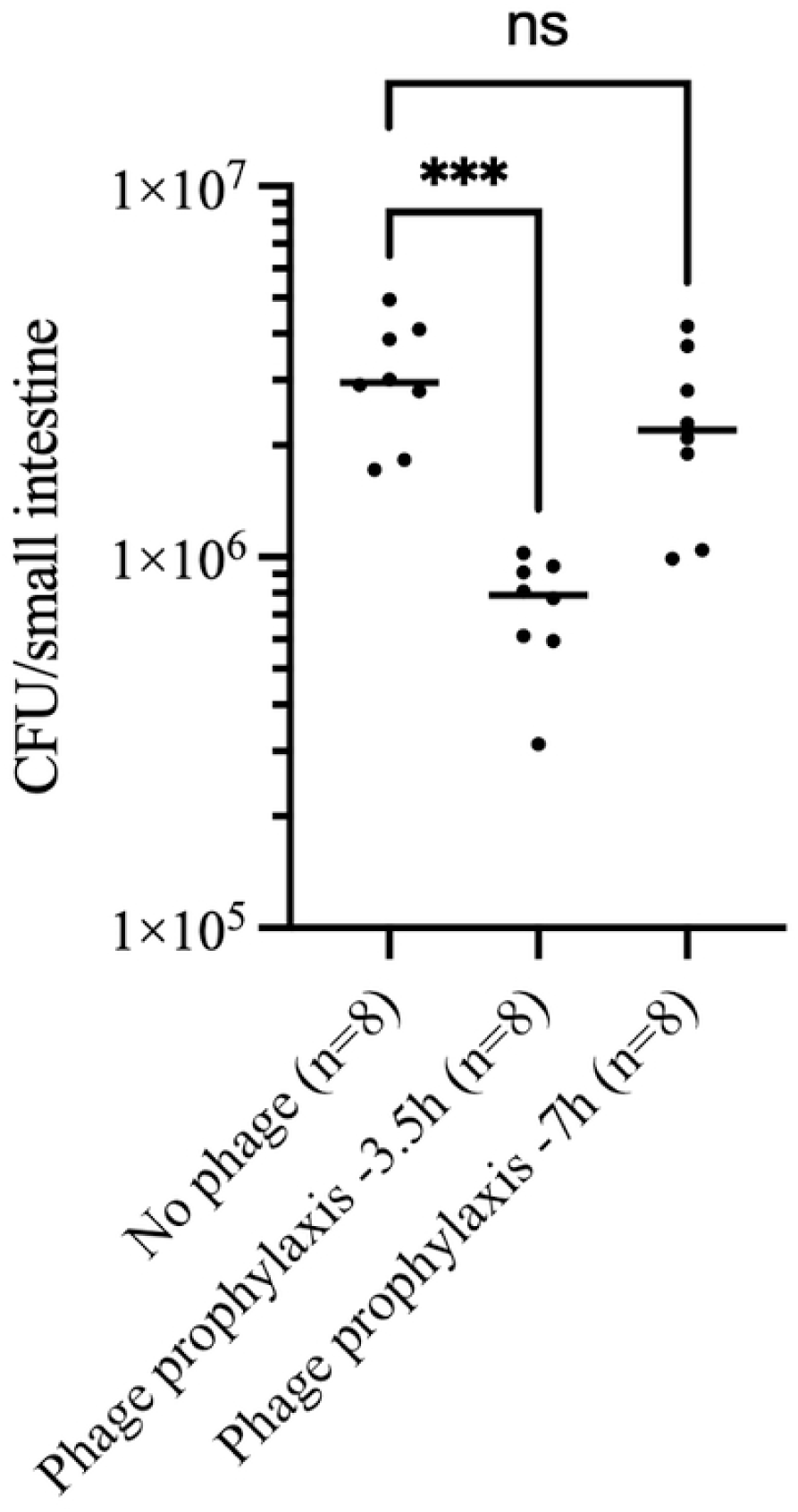
Efficacy of MAA01 phage in reducing *V. cholerae* colonization of the infant mouse small intestine. Mice were intragastrically inoculated with 10^8^ PFU of MAA01 phage 3.5 or 7 h before challenge with 10^5^ CFU *V. cholerae* HC1037. A control group did not receive phage. *V. cholerae* that survived in the small intestine after 21 h were enumerated as CFU per small intestine. The horizontal solid bars represent the medians. Each data point represents one animal. Significance was calculated using the Kruskal–Wallis test with Dunn’s post hoc multiple comparisons test. ***P<0.001.

### Concluding Remarks

In this report, we describe a novel *V. cholerae* O1-specific phage, MAA01, found in both cholera stool samples and aquatic reservoirs in Haiti. MAA01 is phenotypically virulent, despite two genes that predict a temperate lifestyle. We show that MAA01 can prey on *V. cholerae* during infection of the small intestine, making it a potential candidate for phage therapy or prophylaxis. This is the second virulent *V. cholerae* phage identified in Haiti, along with ICP2. The origin of ICP2 seems clear, given its close genetic relatedness to ICP2 isolates in South Asia, where cholera is endemic, and from where cholera was accidentally introduced into Haiti in 2010. In contrast, MAA01’s origin remains unknown. The phage may have been resident prior to 2010, using non-toxigenic *V. cholerae* O1 in the Caribbean Sea as a host, and poised to take advantage of the cholera epidemic. In this scenario, our data suggest that MAA01 jumped into humans in March 2015 on the southern coast of Haiti, then spread to distal sites in subsequent months, with evidence of persistence in clinical samples through 2022. Alternatively, MAA01 may have been introduced into Haiti from elsewhere in the world at that time. Given the renewed interest in using phages, particularly phage cocktails, to treat or prevent infections caused by multidrug-resistant bacteria, identifying phages such as MAA01 that are active during bacterial infection of the human host is important. We propose expanding environmental and clinical surveillance in other cholera-endemic countries to detect additional novel virulent phages.

## MATERIALS AND METHODS

### Isolation of phage from cholera patient stool samples

Stool samples from suspected cholera patients were collected at various CTCs/clinics and transported to our infectious diseases laboratory in Haiti. Each sample was screened for the presence of *V. cholerae* by streaking on thiosulfate–citrate–bile salts–sucrose (TCBS) selective agar plates (Becton, Dickinson and Company). We then serologically tested isolated colonies for *V. cholerae* O1 serogroup and Inaba or Ogawa serotype. Seventy-five stool samples collected between 2014 and 2017 were positive for *V. cholerae* O1 (**Table S1**). Each of the 75 stool samples was subsequently screened for phage plaques in soft agar overlays embedded with *V. cholerae* clinical isolates from Haiti, HC456 and HC1037. Phage plaques were tested by PCR using diagnostic primers specific for phages ICP1, ICP2, or ICP3 (21). Plaques that were negative by PCR were considered to be potential novel phages. Potentially novel phage plaques were plaque-purified three times, and high-titer phage stocks were then prepared as described (21). We subjected eight putative novel phages to whole-genome sequencing using the Illumina platform. All eight phages were genetically identical and represent the novel phage MAA01. We designed and used diagnostic PCR primers for MAA01 to screen all 75 stool samples for this phage.

### Isolation of MAA01 phage from environmental samples

To determine whether MAA01 was present in the environment, we collected 50 mL of surface water from 30 distinct sites across three major regions: Petit-Goave, Jacmel, and Carrefour (**Fig. 1** and **Table S2**). Environmental water sources included: (a) fresh water (flowing river water), (b) wastewater, and (c) estuary water. Of the 30 water samples, 14 were collected from Petitgoave, ten from Jacmel, and six from Carrefour (**Table S2**). After collection, we transported the water samples to our laboratory in Haiti. Each sample was centrifuged at 5000 RCF for 10 min at ambient temperature to pellet particulates. We filtered the supernatant through a 0.22 µm filter (MilliporeSigma) and stored the filtrate at 4 °C. We enriched phages by growing them on two *V. cholera*e O1 strains, HC456 and HC1037 (**Table S2**). Briefly, we used four glass culture tubes per sample; each tube contained 1 mL of 10-fold-concentrated LB broth, 9 mL of potentially phage-containing filtrate, and 100 µL of *V. cholerae* culture (grown to mid-exponential phase). The culture tubes were incubated overnight at 37 °C with aeration. Following incubation, each culture was transferred to a 15 mL conical tube and centrifuged at 5000 RCF for 10 min at 4 °C. The supernatant was filtered through a 0.22 µm filter and stored at 4 °C. The filtered supernatant was serially diluted (10-fold) in LB broth to a final dilution of 10^-5^ in a 96-well microtiter plate, with 90 µL remaining in each well. One hundred microliters of *V. cholerae* culture grown to mid-exponential phase was added to the undiluted culture and to each serial dilution in the 96-well plate, and the plate was incubated at ambient temperature for 10 min to allow phage adsorption. The 190 µL mixtures were subsequently transferred to wells of a six-well plate. Three mL LB soft agar (0.35% agar) cooled to 48 °C was added to each well of the six-well plate, gently swirled to mix the culture and soft agar, and allowed to cool and solidify at ambient temperature for approximately 30 min. The six-well plates were incubated at 37 °C and observed for plaques at 4 and 24 h.

### Whole genome sequencing, genome assembly, and annotation

We selected eight MAA01 phages isolated from different sources for whole-genome sequencing. DNA from each high-titer phage stock was extracted as described previously (21). We constructed the sample library using the Nextera XT DNA Library Preparation Kit (Illumina, Inc.) and sequenced the DNA on the Illumina MiSeq platform for 50 cycles (Illumina, Inc.). We obtained over 200-fold coverage, enabling *de novo* assembly of each phage genome into a single complete contig using CLC Genomics Workbench (Qiagen, Inc.). The GenBank accession number for MAA01 is QB327871.

To identify the receptor for MAA01, we isolated spontaneous MAA01 escape mutants using *V. cholerae* strains E7946 and HC1037. We performed two independent selections for each *V. cholerae* strain and selected three or four clones from each culture for whole-genome sequencing, along with each parent strain. Each genome was assembled *de novo* and analyzed for single-nucleotide polymorphisms (SNPs) and deletion-insertion phenotype (DIP) using CLC Genomics Workbench (Qiagen, Inc.) (**Tables S4 and S5**).

### Phylogenetic analysis

To compare MAA01 phage with existing *V. cholerae*-specific phages and a diverse list of other phages, we constructed a phylogenetic tree based on the amino acid sequence of the Terminase large subunit protein. The MEGA software package was used to construct the phylogenetic tree (35). Briefly, we used the Gonnet substitution matrix to perform multiple and pairwise sequence alignments, and we constructed the phylogenetic tree using the neighbor-joining method (36). We used bootstrap approximation (1000 replicates) to assess the robustness of the internal branches of the neighbor-joining phylogeny.

To support the phylogeny determined using the Terminase large subunit, we used the Virfam bioinformatics tool (34), which identifies and classifies phages based on conserved structural modules. It identifies head, neck, and tail protein modules, as well as recombinase genes, within phage genomes. The platform incorporates a curated database of 1,071 precomputed phage genomes (called ACLAME), enabling comparative analysis.

### Phage adsorption, growth curve, eclipse time, and burst size

For the adsorption experiment, a single colony of *V. cholerae* HC1037 was grown overnight in LB broth at 37 °C with aeration. The culture was diluted in fresh LB broth and grown for approximately 2 hours to reach an optical density OD_600_ of 0.3. Thirty mL of bacterial culture was mixed with 30 µL of MAA01 phage to obtain an MOI of 0.01. The resulting mixture was incubated at ambient temperature for 5 min to allow adsorption; the culture was then washed once with PBS (pH 7.0) and resuspended in 30 mL of LB broth. The culture was incubated at 37 °C with aeration. One mL of culture was collected every 3 min for 21 min. Each aliquot was immediately treated with 1% (v/v) chloroform by vigorous vortexing, then centrifuged at 5000 RCF for 10 min at 4 °C. The supernatant containing phage was then titered by standard plaque assay. Phage plaques were observed at 8 and 24 h after incubation at 37 °C.

To determine the dynamics of MAA01 phage growth, a 30 mL infection culture was prepared as above. We took two 0.5 mL samples at 10 min intervals over 180 min. One sample was mixed with 3 mL of soft agar and plated immediately, while the other sample was plated following treatment with chloroform as described above. The plates were incubated at 37 °C, and phage plaques were observed and counted at 8 and 24 h. A growth curve experiment for MAA01 phage was also performed at 25 °C. Results were plotted to determine eclipse time and burst size. The eclipse time of this phage is 50 min, and the burst size is 95.

### Transmission electron microscopy

High-titer phage stock was subjected to centrifugation at 30,000 х *g* for 1 h in an ultracentrifuge (Beckman Coulter, Inc.). The phage pellet was washed twice with ammonium acetate buffer (0.1 M, pH 7.0). The phage sample was dialyzed in water using a Millipore D-tube dialyzer, MWCo 12-14 kDa (MilliporeSigma, Inc.). We fixed the dialyzed sample with glutaraldehyde, adhered it to a poly-L-lysine-treated, carbon-coated Formvar copper grid, and washed it with PBS. We removed excess solution with filter paper, floated the grid on 1% aqueous uranyl acetate for 30 seconds, and then wiped off excess stain and air-dried the grid. We wiped off excess stain with filter paper and air-dried the grid. We examined phage with an FEI Tecnai G2 Spirit Twin TEM (FEI Corp.) and acquired digital images with a Gatan UltraScan 2k x 2k camera and Digital Micrograph software (Gatan, Inc.).

### MAA01 phage host range determination

Host range was tested by assaying for plaque formation on soft agar overlays using the strains and species listed in **Tables 2** and **S4**. The plates were incubated at 37 °C for 8 and 24 h and then examined for plaques.

### Animal experiment

We performed an infant mouse colonization assay to determine whether MAA01 could reduce *V. cholerae* load in the small intestine. The experiment was conducted at Tufts University under an approved IRB protocol, with methods described previously (27).

Briefly, 5-day-old CD1 mice (both sexes) were anesthetized with 2% isoflurane by inhalation. 10^8^ PFU of MAA01 phage was administered intragastrically via PE10 tubing (Becton, Dickinson and Company) in a volume of 50 µL to pups. We then returned the pups to their dam. After 3.5 or 7 h post-phage administration, we anesthetized the mice again and administered 10^5^ CFU of *V. cholerae* HC1037 intragastrically. Mice were housed without their dam in a 25 °C incubator for 21 h, then euthanized, and their small intestines were removed. We homogenized each small intestine in 1 mL of PBS and plated serial dilutions on LB agar supplemented with 100 µg/mL streptomycin to determine the *V. cholerae* load. Significance was calculated using the Kruskal– Wallis test with Dunn’s post-hoc multiple comparisons test. ***P<0.001.

## ACKNOWLEDGMENTS

This work was supported by National Institutes of Health grants AI097405, AI126357, and AI128750 to JGM, and AI055058 to AC.

